# The Human Nickel Microbiome and its relationship to Allergy and Overweight in Women

**DOI:** 10.1101/546739

**Authors:** E.A. Lusi, I. Santino, A. Petrucca, V. Zollo, F. Magri, D O’Shea, A. Tammaro

**Affiliations:** St Vincent Health Care Group-St Vincent Private Hospital-UCD, Dublin, Ireland; Department of Molecular and Clinical Medicine, School of Medicine and Psychology, Sapienza University of Rome, Microbiology Unit, Sant’Andrea University Hospital, Rome, Italy.; Professional and Allergologic Dermatology Department of Sant’Andrea Hospital, University of Rome La Sapienza, Italy; St Vincent University Hospital-UCD, Immunology and Obesity Research Group, Dublin, Ireland

**Author notes:** Corresponding Author: Elena Angela Lusi (M.D.; Ph.D.).

**Keywords:** Nickel, allergy, women, gut microbiome, overweight

## Abstract

**Introduction:** Nickel exposure usually presents as Allergic Contact Dermatitis. However, Nickel not only causes dermatitis, but an excess of dietary Nickel is reported to be responsible for overweight, metabolic disorders and imbalance of gut microflora.

**Objective:** The aim of study is to expand a preliminary reported evidence of the presence of Nickel-resistant bacteria isolated in human microbiome and further evaluate their association with nickel allergy and overweight in females, the gender mostly affected by Nickel exposure.

**Materials and Methods:** We collected stool samples from 11 lean female with a nickel allergy (BMI <25) and 17 overweight nickel allergic subjects (BMI >25). 11 subjects not allergic to nickel served as control group. Stool cultures were supplemented with increasing concentrations of nickel sulphate (NiSO_4_⋅6H_2_O) from 0.1mM up to 50 mM, in both aerobic and anaerobic conditions *(culturomics*-approach Lusi, 2017). Stool cultures not supplemented with nickel were used as controls. Identification of Nickel resistant bacteria was made by MALDI-TOF technology.

**Results:** In control subjects, 5 mM NiSO4 was the cut off for microbial growth. Conversely, gut bacteria continued to grow at concentration higher than 5 mM in allergic subjects. In particular, Nickel resistant bacteria able to tolerate 32 mM of NiSO4 was detected in 10% of lean allergic and 29% of overweight allergic females. Gut microbes able to grow in at extremely high NiSO4 concentration (50mM) could only be detected in overweight patients with a severe nickel allergy. At increasing NiSO4 concentration, allergic females, especially those with increased BMI, showed a progressive decrease of *Enterobacteriaceae* along with an increased presence of *Lactobacillaceae*, *Bacillaceae* and *Clostridiaceae* compared to control subjects. Major changes in microbial composition were noted at 50 mM of NiSO4 in overweight allergic females.

**Conclusion:** Overweight females with a nickel allergy harbor gut microbes highly resistant to nickel and the role of these bacterial strains must be further elucidated.

## Introduction

The term microbiota is referred to the entire population of microorganisms colonizing the human body. It consists of symbiotic innocuous bacteria and potential pathogens. Microbioma is indeed referred to the genetic composition of the microbiota [1–2]. The microorganisms genes account for more than 3 million, 150 times more than human genes, thus supplementing the host with a “second genome” and contributing to human genetic capacity [3]. Over the last years, the human gut microbiome and its role in both health and disease has been the subject of extensive research. The gut microbiota might be considered as “an individual identity card” which enhances genetic variation among individuals. Over 100 trillion microorganisms colonize the human gut, reaching levels of more than 1000 different species. The most abundant phyla of the adult gut microbiota are Bacteroidetes and Firmicutes [4]. Alterations of gut microbiota are involved in human metabolism, nutrition, physiology, and immune function. Imbalance of the normal gut microbiota have been linked with gastrointestinal conditions such as inflammatory bowel disease (IBD) and irritable bowel syndrome (IBS), and wider systemic manifestations of disease such as obesity, diabetes and atopy [5–8]. Recent evidences suggest that an excess of dietary heavy metals, specifically Nickel, has a profound impact on human health [9–10].

Nickel is the 24th most abundant element on Earth [11–12]. As a natural element of the earth's crust, small amounts of Nickel are found in water, soil, and natural foods. Major dietary source of Nickel is plant food that contain more nickel than animal tissues [13]. While Nickel in small quantities is essential to many biological process, exposure to high concentrations causes skin sensitization. Nickel is an ideal sensitizing agent and it is responsible for the highest incidence of skin sensitization in the industrialized world, even in the paediatric age [14–15]. In sensitized patients, nickel exposure can present either as allergic contact dermatitis (ACD), either as systemic contact dermatitis (SCD). SCD (known as SNAS - Systemic Nickel Allergy Syndrome) follows nickel ingestion and it is characterized by general malaise fever, headache, diarrhea, arthralgia and also eczematous lesions [16].

However, Nickel not only affects the skin, but also causes metabolic abnormalities. In recent times, it has been reported that high contents of dietary Nickel increase the risk of weight gain, especially in women in their menopausal age [17]. The prevalence of Nickel allergy in overweight women is 63%, compared with a prevalence rate of 12% in the general population, and a normocaloric diet formulated to be low only in Nickel is effective in reducing BMI and waist circumference in overweight females. These findings have been recently confirmed in a larger cohort of 1128 obese people, where a Nickel allergy is more frequent in presence of weight excess and is associated with worse metabolic parameters [18].

The excess of Nickel in the diet, not only appears to be a risk factor contributing to obesity, but also causes profound unbalance of the commensal gut flora. Nickel-resistant bacteria are commonly isolated from polluted waters of metallurgic electroplating [19–24], but potentially pathogenic Nickel resistant bacteria have been recently identified also in human gut of obese people [25].

In this study, we aimed to expand preliminary evidences of the presence of nickel resistant bacteria in human gut and study and their correlation with nickel allergy severity and overweight especially in females that are the gender mostly affected by chronic exposure to nickel.

## Materials and Methods

### Sampling

Stool samples from 39 recruited patients, all females, have been collected in the Professional and Allergologic Dermatology department of Sant’Andrea Hospital, University of Rome La Sapienza (Italy) within the period of May-November 2018. The local Institutional Review Board approved the study and all participants provided informed consent.

28 subjects have been diagnosed having Nickel hypersensitivity. The allergic subjects were divided in two subgroups according to their BMI. 17 allergic subjects had a BMI > 25 and 11 allergic had a BMI < 25.

11 subjects not allergic to nickel, (6 overweight and 5 lean), have been recruited as control group. The average age for allergic patients was 44± 3.2 years, while for not allergic controls subjects was 42±4.1 years. For each patient a personal history of allergies, body mass index (BMI) and presence and severity of nickel hypersensitivity were evaluated. Nickel hypersensitivity was determined with a patch test. Subsequently a sample of fresh stool was collected from each enrolled subjects. Within 30 minutes from defecation, the stool samples have been immediately refrigerated at 4°C and processed within 2 hours at the Microbiology Laboratories, Sant’Andrea Hospital, University of Rome La Sapienza.

### Isolation and screening of Nickel resistant bacteria

A sterile solution of Nickel Sulphate 250 mM, prepared by dissolving 66g of NiSO4 ⋅ 6H2O in 1 L of water in a laminar flow hood, has been used to prepare a series of liquid cultures in Brain Heart Infusion (BHI; Becton Dickinson) with increasing concentrations of NiSO4.

Specifically, the concentrations used were 0.1, 0.2, 0.5, 1, 5, 10, 32, 50 mM NiSO4. Stool cultures not supplemented with nickel were used as controls of growth in each round of experiments. Liquid cultures were incubated at 37°C under anaerobic conditions, by covering the liquid culture with 2 mL of paraffin oil. To standardize the initial inoculum, a little quantity of feces has been transferred in liquid phase tampon (Fecal Transwab MWE) containing clary blair transport media.

After the stool sample has been homogenized in the transport media, 100 µL of it have been inoculated in every broth prepared, except the one prepared for the negative control. The liquid cultures have been incubated at 37°C and the bacterial growth has been checked daily for 10 days. The liquid cultures showing bacterial growth in presence of NiSO4 have been sub cultivated over agar blood plates and incubated in aerobic and anaerobic conditions.

### Identification of the bacterial isolates

The microbiological identification of the isolated bacteria has been carried out by MALDI-TOF technology. The system used, the Bruker Daltonik IVD MALDI Biotyper 2.3, is capable of comparing and evaluating similar mixtures of complex molecules, extracted from an unknown microorganism, with the database of the instrument. At the end of the procedure, the microorganism is identified for genus and species.

## Results

The research conducted in 39 female showed how different Nickel concentration changes intestinal microbial growth and composition. In control subjects, 5 mM of NiSO4 was the cut off limit of bacterial growth, as bacteria no longer grew at higher concentrations (Figure 1).

**Figure 1.**
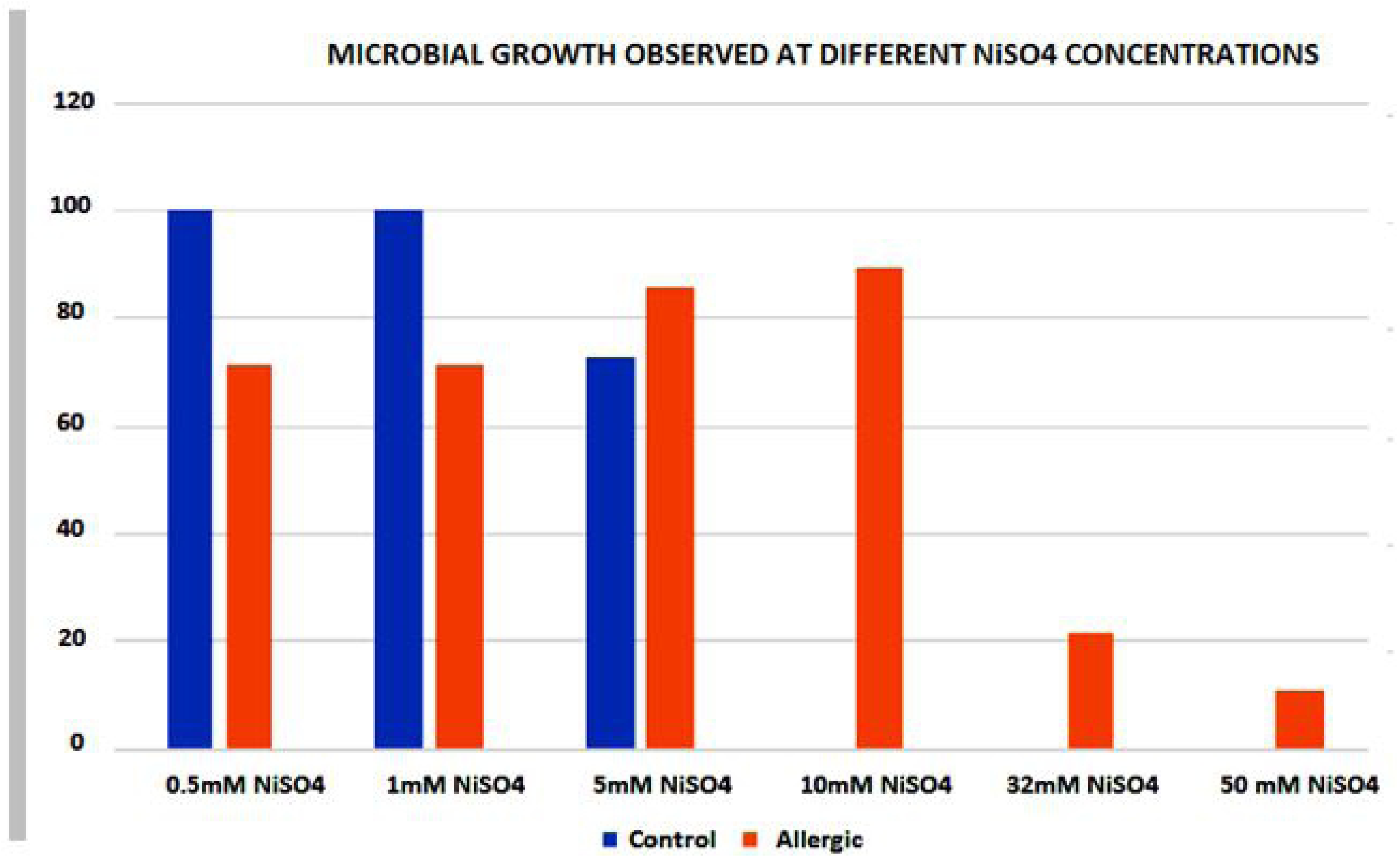
Growth pattern of Nickel resistant bacteria in human gut of control subjects versus allergic patients with a Nickel allergy.

While for control subjects the cut off is 5 mM NiSO4, in the allergic patients it is possible to observe how gut bacteria continue to grow at concentration higher than 5 mM. Allergic patients with a BMI <25 (11 patients) show an intestinal microbiota capable of tolerating lower NiSO4 concentrations, compared to allergic patients with increased BMI. Bacteria stop their growth at 10 mM NiSO4 in the almost of the totality of allergic lean patients and only 10% of them harbor gut microbes able to grow at 32 mM concentration of NiSO4. Conversely, at 50mM NiSO4, that is an extremely high nickel concentration, bacterial growth can be detected only in allergic subject with increased BMI and high severity of nickel allergy at patch test (3+) (Figure 2).

**Figure 2.**
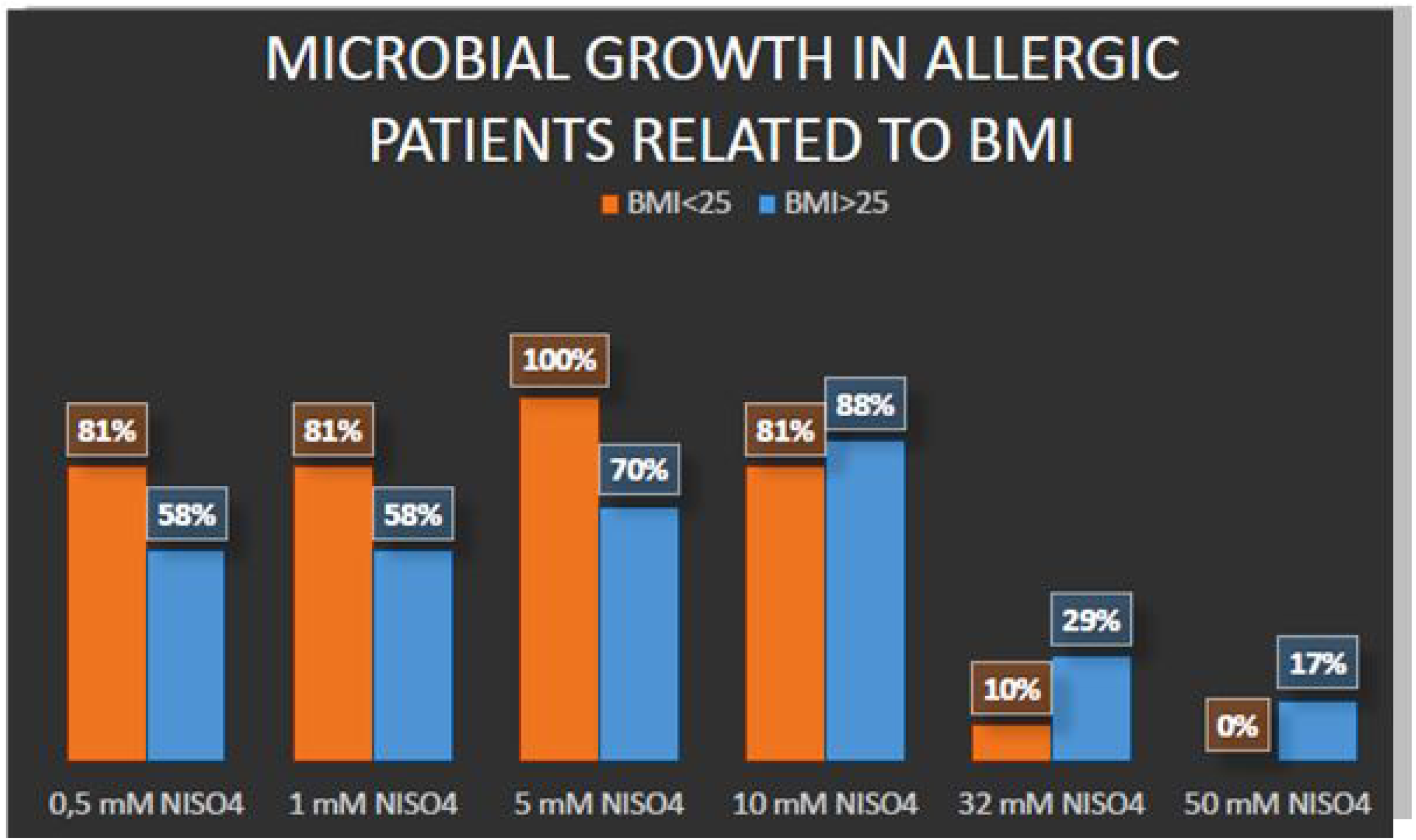
Allergic patienst with BMI >25 harbour gut microbes able to tollerate higher concentration of NiSO4 compared to lean allergic subjects. Gut microbes able to grow in the presence of 50 mM NiSO4 are detected only in overweight allergic patients.

When analyzing the microbial composition, it is possible to observe how microbiota of control subjects is more abundant in *Enterobacteriaceae* compared to allergic patients. In particular, levels of *Enterobacteriaceae* tend to decrease progressively with increasing NiSO4 concentrations, while Lactic Acid Bacteria (*Enterococcaceae*, *Streptococcaceae* and *Lactobacillaceae*) increase. When analyzing the bacterial composition according to bacterial order, *Lactobacillales* are found in higher amount in allergic patients, especially in those with increased BMI and high degree of Nickel allergy severity. Gut microbes able to grow at extremely high NiSO4 concentration (50mM) can be isolated only from overweight allergic patients. At 50 mM NiSO4 changes of gut microbiota become profound with a major reduction of *Enterobacteriales* and increased level of *Bacillales* and *Clostridiales* (Figure 3).

**Figure 3.**
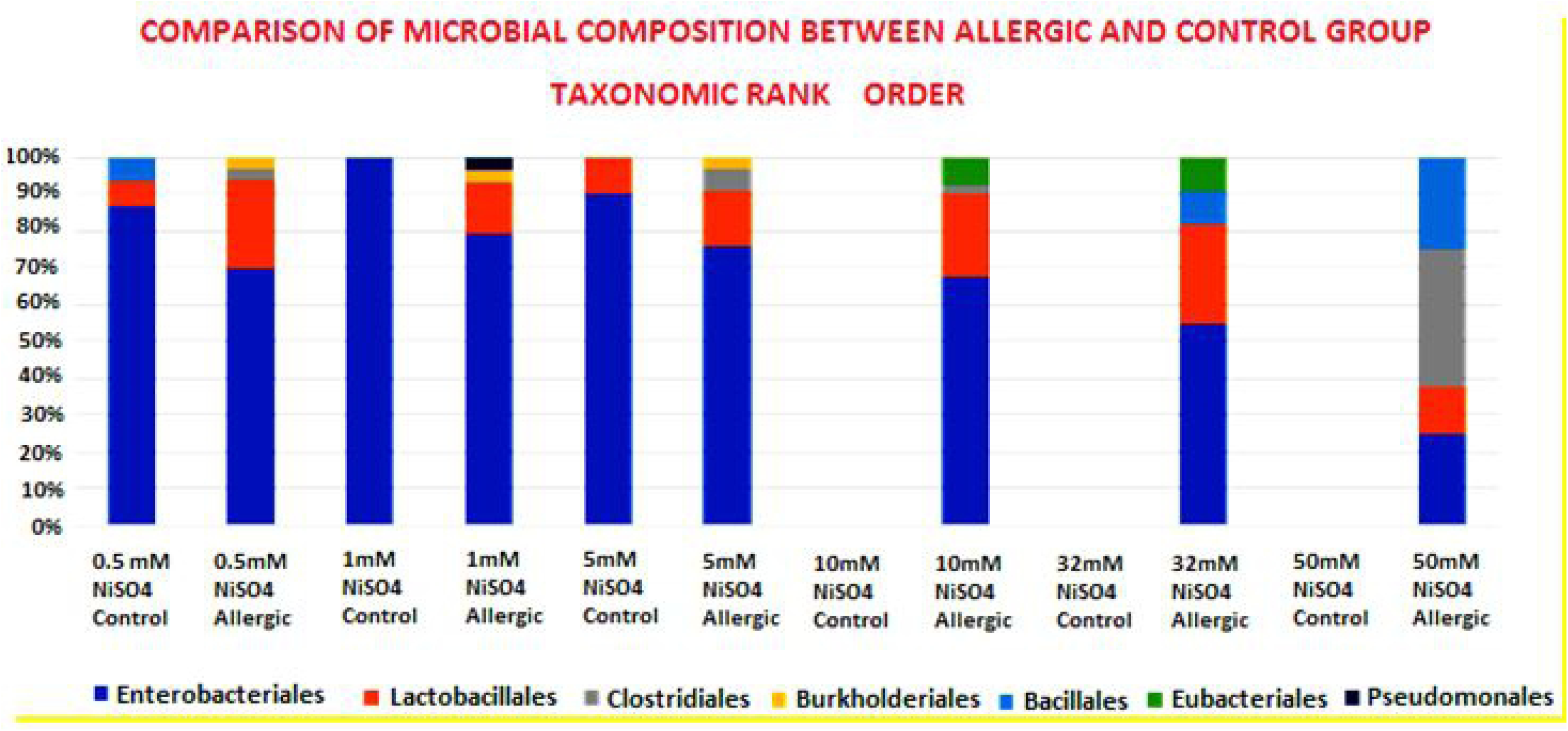
Allergic patients show a decreased level of Enterobacteriales and incresed presence of Lactobacillales compared to control subjects. With increasing NiSO4 concentration, levels of Enterobacteriales tend to decrease progressively while Lactobacillales increase. At 50 mM NiSO4, microbial growth is detected only in allergic overweight patients with a dramatic reduction of Enterobacteriales and increased level of Bacillales and Clostridiales.

## Discussion

In a study conducted in 2015, Lusi et al. described the role of dietary nickel in determining overweight in allergic females in their perimenopausal age [17]. This finding highlighted that Nickel was not only implicated in skin allergy but for the first time an excess of Nickel in the diet was linked to overweight. A normocaloric diet, formulated to be low only in Nickel reduced the BMI and waist circumference in the treated subjects. The fact that an excess of Nickel could be associated to obesity and metabolic abnormalities was confirmed subsequently in a large cohort study [18].

There are numerous animal studies showing that gut microbes can induce obesity [26–31] and Lusi first posed the question if a high dietary Nickel load could also influence the intestinal microbiome and finally demonstrated the presence of Nickel resistant bacteria in the intestinal tracts of overweight subjects [25].

The presence of Nickel resistant bacteria in humans is a recent finding. So far heavy metals resistant microorganisms have been isolated in contaminated mining soils and effluents of industrial sources. [19–24].

Following the same *culturomics* technique, we aimed to replicate and expand these initial findings analyzing the microbial intestinal composition at different concentrations of Nickel Sulphate (NiSO4) in a group of 28 Italian females with a Nickel allergy and 11 controls.

Nickel resistant bacteria were isolated in both allergic and control subjects. However, while in control subjects bacteria stop their grow at 5 mM NiSO4, bacteria isolated from allergic patients were able to tolerate higher concentrations of NiSO4, up to 50mM in obese allergic.

In contrast to the previous paper, where the cut off limit of bacterial growth in allergic obese patients was 1mM NiSO4 [25], in this new round of experiments we could detect gut bacteria able to grow at 50mM concentration of NiSO4 in allergic obese.

It is important to note that 50mM is fraction of the Nickel solution that is commonly used for metallurgic electroplating and never in humans it has been reported bacteria able to tolerate NiSO4 concentration of this magnitude, not even Helicobacter Pylori [32].

With increasing Nickel concentration, microbiota of allergic patients, especially those with increased BMI, becomes less abundant in *Enterobacteriaceae* and increases in “Lactic acid bacteria” (*Enterococcaceae*, *Streptococcaceae* and *Lactobacillaceae* family, comprised by the order of *Lactobacillales*). At 50 mM of NiSO4 the microbiota composition of allergic overweight patients shows a profound dysregulation and a drastic reduction of *Enterobacteriaceae* with increased levels of *Bacillaceae* and *Clostridiaceae*.

In agriculture and mining science, Nickel resistant bacteria have been considered as useful scavengers evolved to eliminate the excess of heavy metals from the environment [33]. Paralleling the function of Nickel resistant bacteria found in soil, it may be that human Nickel bacteria evolved pathways to regulate metal ion accumulation and avoid heavy metal toxicity in human?

Some strains of *Lactobacillales* (e.g. *Lactobacillus fermantum*) use Nickel ingested with diet in order to protect the body from Nickel toxic potential. Lactic acid bacteria bind to toxic substances, such as aflatoxin B_1_ and food-borne mutagen Trp-P-2 within the gastro-intestinal tract, thereby reducing their uptake [34–36]. This might support the hypothesis that Lactic acid bacteria are involved in Nickel detoxification, but this finding might also fuel the controversy about the role of *Lactobacillales* in obesity [37–41].

In this framework is also plausible to think that obese subjects could be populated by particular strains of *Lactobacillales* rather than others and some strains of Nickel resistant *Lactobacillae* could be actually responsible of shaping the obese phenotype.

This study, although simple and technically basic, confirms how an excess of Nickel can influence the gut microbiota in obese Nickel allergic females.

Only additional experiments and the inoculation of human Nickel resistant bacteria in animal models will clarify the role of Nickel resistant bacteria and if the human–Nickel-microbiota is a friend or foe.

## Author contribution

EAL : Concept, description of human nickel resistant bacteria, Study design, culturomics protocol, wrote and edited the manuscript

IS: Microbiology Lab procedures, data analyses and graphical visualization

A.P: Microbiology Lab procedures, data analyses and graphical visualization

ZV : Clinical data collection, logistic

MF : Clinical data collection, logistic

DOS: Study design, study coordinator and critical analyses

AT: Study coordinator, patients enrollment, clinical data analyses, wrote the initial draft of manuscript.

## Conflict of Interest

None to declare.

## Funding Statements

The Authors did not receive specific funding for this work.

